# opentsv prevents the corruption of scientific data by Excel

**DOI:** 10.1101/497370

**Authors:** Peter De Rijk, Svenn D’Hert, Mojca Strazisar

## Abstract

Microsoft Excel is widely used by researchers to edit tab- or comma-separated data files. However, Excel often corrupts the data when opening these files, most notably by changing some gene names to a date. Although this problem was cautioned against earlier, we show that every year hundreds of published papers still come with supplementary data files containing these errors. Opentsv was developed to effectively circumvent this problem at the root by providing an easy and transparent way to open delimited data files in Excel without these conversions. Opentsv is freely available at https://github.com/derijkp/opentsv.

## Background

Bioinformatics analysis software often produces results in tab-separated value (tsv) or comma-separated value (csv) format. For viewing, annotating, editing or formatting these data files, biologists commonly use Microsoft Excel (Microsoft Corp., Redmond, WA, USA) because it provides an easy to use interface to tabular data they are already familiar with. Also, for many journals it is the only accepted file type that can contain tabular data with formatting.

However, already more than a decade ago researchers discovered a serious flaw in Excel for this type of use (Zeeberg et al. 2004); When developing their software for data mining published gene expression data sets (Zeeberg et al. 2003) they kept coming across unknown gene names: They found that these were caused by erroneous conversions by Excel: Gene names that look like dates were transformed to an internal date format when opening tsv or csv data files. Other identifiers such as the RIKEN “2310009E13” were also found to be converted irreversibly to the floating-point number “2.31E+13”.

Similarly, in non-biological fields people are reporting conversion errors. Leading zeros (e.g. in zip-codes and telephone numbers) or trailing zeros (e.g. in version numbers such as 1.10) can be lost in a conversion to a number. Any identifier consisting of numbers and hyphens or dots (e.g. size ranges, product codes, hyphenated room numbers, storage location codes) risks conversion to a date. Ironically, not even dates are completely safe: An unambiguous iso 8601 conformant date such as 2018-02-04 is converted to an ambiguous, locale dependent format (2-4-2018 or 4-2-2018) upon export (Kosmala, 2016). A date where only the month and year are indicated (e.g. april 2001) will be changed to apr-01, which can, after export, be easily confused with the first of April instead.

Zeeberg et al (2004) already expressed the hope that these problems would be solved by making the automatic conversions a non-default option in Excel. However, after more than a decade, there still is no such option.

A more recent screening (Ziemann et al. 2016) of supplementary files accompanying articles and Excel files deposited in the NCBI Gene Expression Omnibus showed that gene name errors continued to be a problem in supplementary files: In different journals tested, from 5% to over 30% of papers with supplementary gene lists in Excel files have converted gene symbols; 39.7% of genelists containing Excel files deposited to NCBI GEO (Barrett et al. 2013) were found to contain gene name errors.

Recently some tools were developed to mitigate the problem: Escape Excel converts the delimited file into an escaped format before loading in Excel (Welsh et al. 2017), while Truke (Mallona et al. 2017) detects and fixes misconverted gene symbols to reduce the damage after the fact. We wanted a transparent fix for the problem at the root, avoiding (mis)conversions all together.

## Results

In order to get an idea of how widespread the problem still is, we screened 15 journals publishing genomics and transcriptomics data and NCBI GEO for supplementary Excel files with corrupted gene lists (Figure 1) using an approach similar to the one used in Ziemann et al. 2016. While the rising trend seen in the Zieman paper seems to have stopped in 2016, it remains at a high level with a total of 616 publications with error-containing supplements detected in 2018. Also in 2018, up to 36% of papers with gene lists in the supplements was affected in the different journals. In GEO 44.6% of submissions with Excel gene lists contained errors.

**Figure 1.**
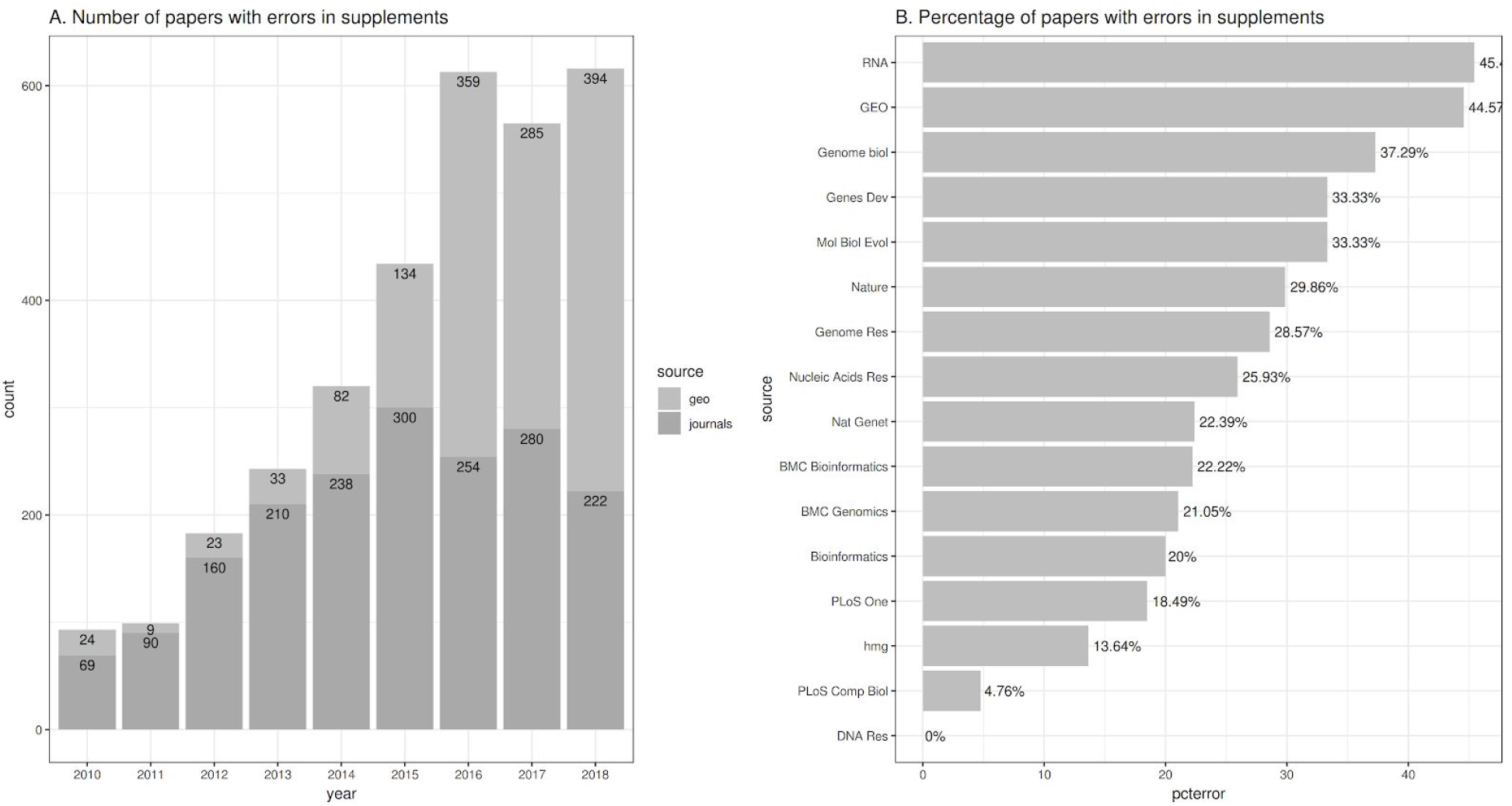
Number and percentages of publications with corrupted Excel files. Panel A shows, per year, the total number of articles where converted gene names were detected in the supplementary files of 15 journals, and the total number of GEO submissions with errors. The numbers give a lower bound as only Excel files detected to contain gene list were scanned for gene name errors. Panel B shows the percentage of publications with a gene list in the supplementary files where erroneous conversions were detected, for journals published in 2018.

Cautioning researchers to carefully handle the data using cumbersome workarounds, or not to use Excel at all, was clearly not sufficient to resolve this problem. Opentsv was created to basically provide Excel users the option to open tsv and csv files safely by default. It is a small program that supports, by simply double clicking, opening tsv and csv files in Excel without automatic conversions. Data files loaded via opentsv are by default (after confirmation) saved as tsv files. This is useful in situations where tsv files can be downloaded, edited and uploaded to a server for bulk changes of tabular data. With opentsv, Excel can be safely used for editing these types of files.

Opentsv is available as a portable executable that can be used with or without installation. Running the opentsv executable directly displays a dialog with some help on the software and several options, including installation (Figure 2). The Install button will copy the executable into the program files folder and link it to the appropriate file types. Alternatively, the “Register opentsv as default program for opening” button can be used to link file types to the program at its current location, allowing its easy use without needing administrator access to the program files folder. Opentsv can be linked to other file extensions as well using the default Windows methods.

**Figure 2.**
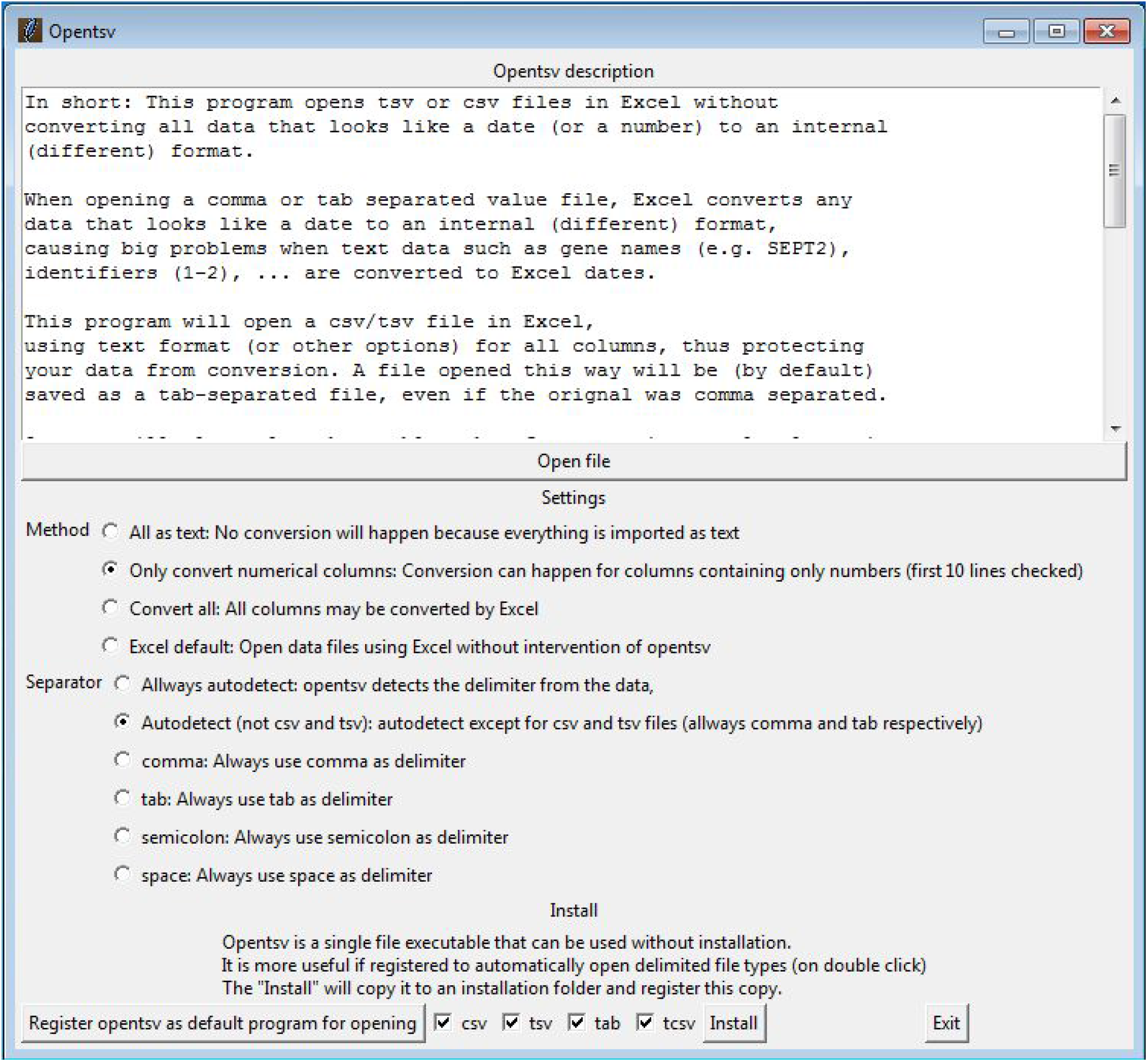
Settings and installation screen

When linked to the file types, double clicking will open the file using opentsv, which subsequently opens Excel and uses the Excel programmatic interface to open the file without the erroneous conversions.

The “Open file” button will allow the user to select and open a file in Excel directly from opentsv using the current settings, which are shown below it. The following options for handling values are available:

- The “**All as text**” option will cause all fields to be loaded as text fields. In this case no conversion happens at all. In order to minimize the chance of erroneous conversion, this is the default.
- In the default settings, numbers are imported as text, and to use them in calculations they have to be manually converted after loading. The “**Only convert numerical columns**” option provides will automatically convert some numbers: If the header field is considered safe from conversion and the first 1000 data lines are all strictly numbers or empty, generic conversion for that column is allowed.
- “**Convert everything**” uses the generic conversion for all fields, turning of protection against misconversions, but still correcting for sepators.
- Finally “**Excel default**” can be used to return (temporarily) to Excel opening data files without intervention of opentsv.

Another Excel problem tackled by opentsv is the use of separators: In some locales (where the comma is used as a decimal separator) Excel diverges from the general csv format specification (https://tools.ietf.org/html/rfc4180) by using semicolons as separator instead of commas. Because of this, a user in such locale cannot easily open a properly formatted csv file (using commas) produced by software or international colleagues: Opentsv supports several options for the use of separator:

- By default (**Autodetect**), it will auto detect the separator from the data except for files with the extension csv and tsv, where respectively the comma and the tab character are used.
- With the “**Allways auto detect**” option, the separator is also autodetected for csv and tsv files.
- The separator can be manually set to **comma, tab, semicolon** or **space**.

Unfortunately the way Excel opens files with the extension .csv seems to be hardcoded in the program and cannot be changed with the APIs opentsv uses, requiring a hack: A .csv file is first copied to a file with the extension .tcsv and subsequently opened.

## Conclusions

We show that the auto-conversion of values when loading comma or tab delimited data files in Excel continues to lead to erroneous data in published literature and databases. Most researchers however are not willing to forego the conveniences of Excel. For more than a decade users have, without success, been asking Microsoft for an option to turn of this conversion. Opentsv finally provides this option in an easy to install program that intervenes mostly transparently in loading data into Excel without the conversion. Its widespread use may help to reduce the significant number of these errors in published literature.

## Methods

The script used to scan 15 journals and GEO for errors in gene lists created by Excel conversion is available at the opentsv github site (https://github.com/derijkp/opentsv/tree/master/checkpapers). The script uses genomecomb shell (Reumers et al. 2011) to run. Urls for all papers between 2010 and 2019 were obtained by either scraping the journal website directly using wget or by querying pubmed, saving the result as xml and parsing the result. For PLoS One this query was limited to only extract papers with genome or transcriptome in the abstract. Entries for GEO were obtained by querying in the web interface and parsing the resulting files. The papers were downloaded and parsed for links to Excel supplementary files. The Excel files were downloaded and converted to text using ssconvert. Detection of gene lists and errors in these gene names was implemented using the same algorithm as used in Ziemann et al. (2016).

Opentsv is implemented in Tcl and made into a portable, single file executable using tclkit. Source code as well as executables are available at https://github.com/derijkp/opentsv under the MIT license.

## Declarations

### Ethics approval and consent to participate

Not applicable

### Consent for publication

Not applicable

### Availability of data and material

Opentsv is freely available at https://github.com/derijkp/opentsv. The source code is available under the MIT license. The code used for scanning the journals is available with the source.

### Authors’ contributions

PDR designed the study with input from MS and SD. PDR performed the analysis and wrote the software, with testing and feedback from MS and SD. PDR wrote the manuscript together with MS and SD. All authors read and approved the final manuscript.

## Acknowledgments

Not applicable

